# An AI-assisted Tool For Efficient Prostate Cancer Diagnosis

**DOI:** 10.1101/2022.02.06.479283

**Authors:** Mustafa Umit Oner, Mei Ying Ng, Danilo Medina Giron, Cecilia Ee Chen Xi, Louis Ang Yuan Xiang, Malay Singh, Weimiao Yu, Wing-Kin Sung, Chin Fong Wong, Hwee Kuan Lee

## Abstract

Pathologists diagnose prostate cancer by core needle biopsy. For low-grade and low-volume cases, the pathologists look for the few malignant glands out of hundreds within a core. They may miss the few malignant glands, resulting in repeat biopsies or missed therapeutic opportunities. This study developed a multi-resolution deep learning pipeline detecting malignant glands in core needle biopsies to help pathologists effectively and accurately diagnose prostate cancer in low-grade and low-volume cases. The pipeline consisted of two stages: the gland segmentation model detected the glands within the sections and the multi-resolution model classified each detected gland into benign vs. malignant. Analyzing a gland at multiple resolutions provided the classification model to exploit both morphology information (of nuclei and glands) and neighborhood information (for architectural patterns), important in prostate gland classification. We developed and tested our pipeline on the slides of a local cohort of 99 patients in Singapore. The images were made publicly available, becoming the first digital histopathology dataset of prostatic carcinoma patients of Asian ancestry. Our pipeline successfully classified the core needle biopsy parts (81 parts: 50 benign and 31 malignant) into benign vs. malignant. It achieved an AUROC value of 0.997 (95% CI: 0.987 - 1.000). Moreover, it produced heatmaps highlighting the malignancy of each gland in core needle biopsies. Hence, our pipeline can effectively assist pathologists in core needle biopsy analysis.

## 1 Introduction

Prostate cancer is the second most common cancer diagnosed in men worldwide [1]. It is diagnosed by core needle biopsy analysis, involving a collection of about 12 cores from different parts of the prostate. Pathologists analyze individual prostate glands on the slides of the collected cores for malignancy. For low-grade and low-volume cases, pathologists have to carefully examine hundreds of glands in each core to avoid missing any malignant glands. This is a tedious and time-consuming process, prone to errors and inter-observer variability. Besides, increasing incident rates and decreasing number of actively working pathologists escalate the workload per pathologist [2].

Prostatic carcinoma is graded based on Gleason patterns (GPs) from 1 to 5 (although GP1 and GP2 are not routinely reported in the clinic). The sum of the two most common patterns inside the slide (for biopsy slides, the highest grade is reported as the second pattern if it is > 5%) is the Gleason score (GS) and used as the grade of prostatic carcinoma. Recently, the 2014 International Society of Urological Pathology (ISUP) Consensus Conference on Gleason Grading of Prostatic Carcinoma has updated the definitions of GPs and introduced a new grading system [3]. There are five prognostically different grade groups from GG1 to GG5 (from the most favorable to the least favorable), based on the modified GS groups: GG1 (GS ≤ 6), GG2 (GS 3 + 4 = 7), GG3 (GS 4 + 3 = 7), GG4 (GS 8), and GG5 (GS 9 — 10). It is vital to diagnose cancer at low grades for a better prognosis and patient life quality.

Recently, it has been shown that the assistance of machine learning systems significantly improves the diagnosis and grading of prostate cancer by pathologists [4, 5]. A few studies developed successful machine learning systems for prostate cancer diagnosis and grading to assist pathologists [6–14]. They covered a broad spectrum of grade groups. Nevertheless, it is easier for an expert pathologist to diagnose high-grade cancers such as GG4 or GG5. On the other hand, it becomes a real challenge to discriminate rare malignant glands among numerous benign glands in low-grade cancers such as GG1 and GG2 [15]. Besides, undetected malignant glands may result in repeat biopsies and missed therapeutic opportunities. Therefore, this study concentrates on low-grade prostatic carcinoma of GG1 and GG2. It develops a deep learning pipeline detecting rare malignant glands in core needle biopsy slides to help pathologists quickly and accurately diagnose prostate cancer in low-grade and low-volume cases. Given a core needle biopsy slide, the pipeline produces a vivid heatmap highlighting the malignant glands inside the slide to the pathologists. Moreover, contrary to previous studies’ single resolution and patch-based methods, this study uses a multi-resolution and gland-based classification approach, providing pathologists with individual gland labels.

Our pipeline consists of two stages: gland segmentation using a Mask R-CNN [16] model and a novel multi-resolution gland classification model (see Methods 2.2 and Figure 1). While glands in biopsy cores were detected by the gland segmentation model, each detected gland was classified into benign vs. malignant by the gland classification model. Our multiresolution gland classification model jointly analyzed a gland’s high resolution (40 × and 20×) and low resolution (10× and 5×) patches to exploit morphology information (of nuclei and glands) and neighborhood information (for architectural patterns) (see Methods 2.2.2), respectively. The multi-resolution models imitate the pathologists’ comprehensive workflow of analyzing both macro and micro structures inside the slides [17, 18], and even naive multiresolution models obtained as ensembles perform better than single-resolution models [6]. The code is publicly available at Zenodo under the https://doi.org/10.5281/zenodo.5982398.

**Figure 1:**
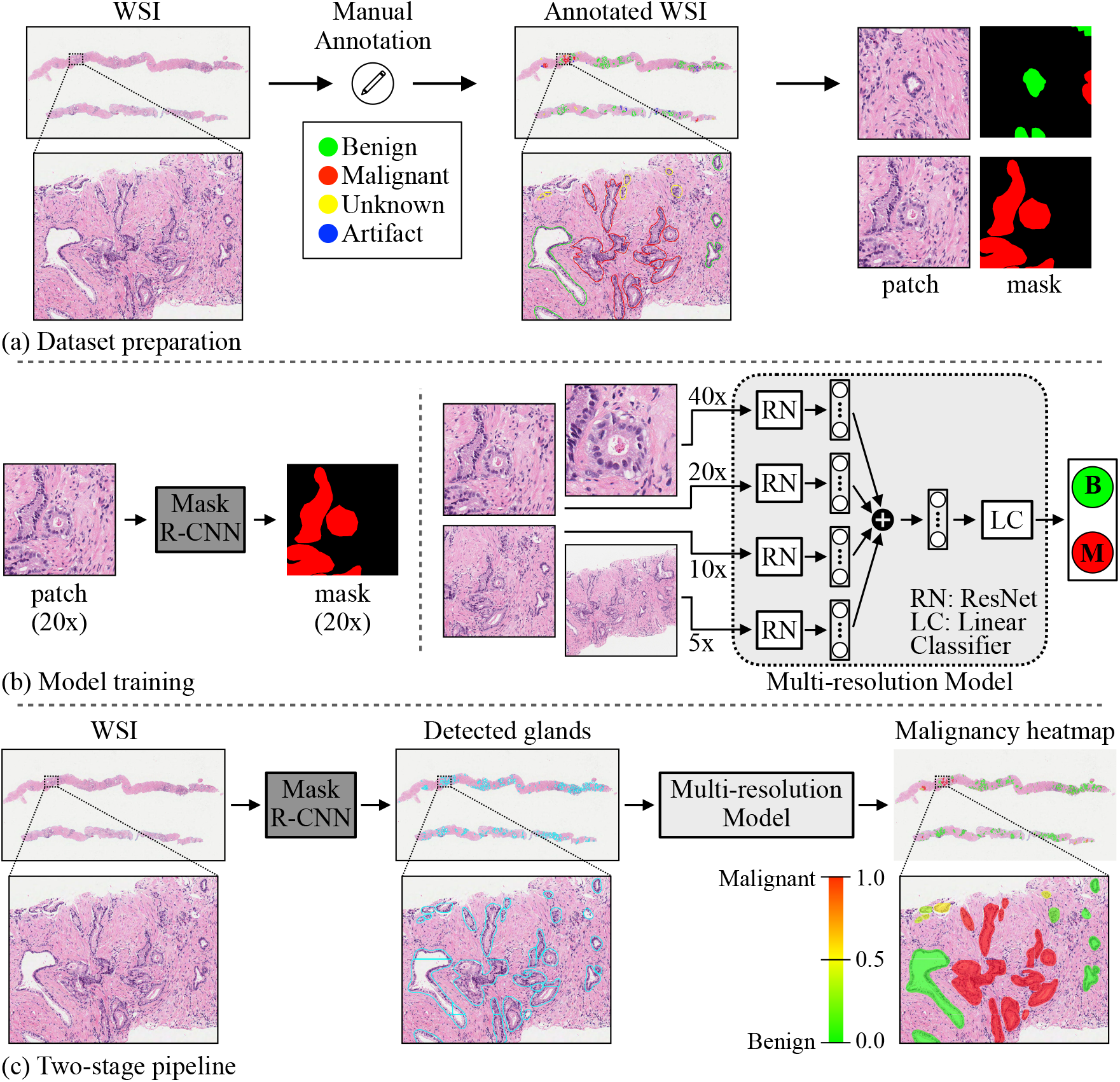
Malignant gland detection pipeline in prostate core needle biopsies. (a) Glands in collected WSIs are annotated by drawing contours around each gland and assigning a malignant, benign, unknown or artifact label. Then, patches centered around a particular gland are cropped at different resolutions for each gland while preparing datasets for training machine learning models. (b) A Mask R-CNN [16] model is trained for gland segmentation on patch and mask pairs prepared at 20× resolution. A multi-resolution (four-resolution) deep learning model is trained for gland classification on patches (cropped at 5×, 10×, 20×, and 40× resolutions) and label pairs. (c) After gland segmentation and classification models are successfully trained, a two-stage pipeline is constructed. The trained Mask R-CNN model detects the glands in a WSI, and the trained multi-resolution model obtains the malignancy probability for each detected gland. Finally, a malignancy heatmap is generated to support pathologists in prostate cancer diagnosis from core needle biopsy WSIs.

This study was conducted on a local cohort of 99 patients collected in Singapore (see Methods 2.1 and Table 1). Data collected from each patient contained hundreds of glands for machine learning. The data is publicly available [19], and this is the first digital histopathology dataset of prostatic carcinoma patients of Asian ancestry. The global research community can benefit from this valuable dataset to develop and test machine learning models and ultimately improve patients’ outcomes.

**Table 1:** The number of slides and patches in training, validation, and test sets for gland segmentation and classification tasks. There is one H&E stained WSI for each prostatectomy or core needle biopsy specimen. The gland classification datasets are the subsets of the gland segmentation datasets.

| Gland Segmentation |  |  |  |  |  |  |  |  |
| --- | --- | --- | --- | --- | --- | --- | --- | --- |
|  | # slides |  |  |  | # patches |  |  |  |
|  | train | valid | test | total | train | valid | test | total |
| prostatectomy | 17 | 8 | 15 | 40 | 8545 | 4014 | 7543 | 20102 |
| biopsy | 26 | 13 | 20 | 59 | 5858 | 4419 | 6737 | 17014 |
| total | 43 | 21 | 35 | 99 | 14403 | 8433 | 14280 | 37116 |
| Gland Classification |  |  |  |  |  |  |  |  |
|  | # slides (3+3:3+4:4+3) |  |  |  | # patches (benign:malignant) |  |  |  |
|  | train | valid | test | total | train | valid | test | total |
| biopsy | 10:9:1 | 3:7:0 | 6:10:0 | 19:26:1 | 1557:2277 | 1216:1341 | 1543:2718 | 4316:6336 |

The pipeline’s performance was evaluated on the data of held-out patients in the test set. The performance metric was the area under the receiver operating characteristic curve (AUROC) value with a 95% confidence interval (CI) constructed using the percentile bootstrap method [20]. We obtained an AUROC value of 0.997 (95% CI: 0.987 - 1.000) on benign vs. malignant classification of core needle biopsy parts (81 parts: 50 benign and 31 malignant) in 16 slides of 16 patients in the test set. Furthermore, we produced spatial malignancy maps with a gland-level resolution to assist pathologists in reading prostate core needle biopsy slides (Figure 3). Our pipeline can help pathologists detect prostate cancer in core needle biopsy slides at early stages and shorten turnaround times by presenting high-risk regions via malignancy maps to the pathologists.

## 2 Materials and Methods

### 2.1 Dataset

Digitized haematoxylin and eosin (H&E)-stained whole-slide-images (WSIs) of 40 prostatectomy and 59 core needle biopsy specimens were collected from 99 prostate cancer patients at Tan Tock Seng Hospital, Singapore. There were 99 WSIs in total such that each specimen had one WSI. H&E-stained slides were scanned at 40× magnification (specimen-level pixel size 0.25*μm* × 0.25*μm*) using Aperio AT2 Slide Scanner (Leica Biosystems). Institutional board review from the hospital were obtained for this study, and all the data were de-identified.

We developed models for the gland segmentation and gland classification tasks using the 99 WSIs. The 99 WSIs were randomly segregated into training (43), validation (21), and test sets (35) at the patient level to avoid data leakage while training the models [21]. While all the slides were utilized in the gland segmentation task, only a subset of the slides in each set (training: 20, validation: 10, and test: 16) was used in the gland classification task (Table 1). The models were trained on the training sets. The best sets of model weights were chosen on the validation sets using early stopping to avoid overfitting, and the best models were evaluated on the test sets.

Prostate glandular structures in core needle biopsy slides were manually annotated and classified into four classes: benign, malignant, unknown, and artifact (Figure 1a), using the ASAP annotation tool (https://computationalpathologygroup.github.io/ASAP/). A senior pathologist reviewed 10% of the annotations in each slide, ensuring that some reference annotations were provided to the researcher at different regions of the core (see Appendix A for details). It is to be noted that partial glands appearing at the edges of the biopsy cores were not annotated.

### 2.2 Prostate gland segmentation and classification pipeline

This study developed a deep learning-based pipeline detecting malignant glands in core needle biopsy slides of prostate tumors. The ultimate aim was twofold: to improve patients’ outcomes by helping pathologists diagnose prostate cancer in low-grade and low-volume cases and to reduce the pathologists’ workload by providing them with an assistive tool during diagnosis. The pipeline consisted of two stages: gland segmentation and gland classification models.

#### 2.2.1 Gland segmentation using a Mask R-CNN model

The first stage used a Mask R-CNN [16] model to segment glands. The dataset used in this stage consisted of cropped patches of size 512 × 512 pixels at 20× magnification from whole slide images such that an annotated gland was centered at each patch (Figure 1b). The patch size and resolution were selected such that both nuclei morphology and gland structure information were available to be exploited by the Mask R-CNN model. For each patch, binary masks of all glands present in the patch, including incomplete glands at the edges, were created as labels.

Data augmentation techniques, namely random horizontal and vertical flip, colour augmentation (contrast, brightness), and rotation, were applied to the patches and binary masks at training. After augmentation, the patches were cropped to a size of 362 × 362 pixels around the centre and passed as input to Mask R-CNN.

#### 2.2.2 Gland classification using a four-resolution model

The second stage used a four-resolution deep learning model that emulates pathologists’ workflow to perform gland classification. Patches of size 512 × 512 pixels were cropped from whole slide images at resolutions 5×, 10×, 20×, and 40× with an annotated gland centered at each patch. To predict whether the center gland was benign or malignant, patches of these resolutions from the same tissue region (around a particular gland) were passed into the multi-resolution model simultaneously (Figure 1b).

Specifically, each patch of a different resolution was passed to a different ResNet-18 [22] feature extractor. Extracted features from patches of all resolutions were then summed and passed to a linear classifier to predict whether the center gland was benign or malignant. The same data augmentation techniques used in the first stage were applied during the training of the multi-resolution model. The model was trained end-to-end.

### 2.3 Training of deep learning models

The Mask R-CNN model was trained using the Adam optimizer with a batch size of 4 for 142 epochs. The learning rate was initially set to 3e-4. After the training loss plateaued at the end of epochs 60 and 110, it was reduced to 3e-5 and 3e-6, respectively. Similarly, the multi-resolution model was trained using the Adam optimizer with a learning rate of 5e-4 and a batch size of 32 for 76 epochs.

### 2.4 Inference using trained pipeline

The trained pipeline accepted a whole-slide image of prostate core needle biopsy as input, detected the glands within the slide, and finally predicted whether each detected gland was malignant or benign (Figure 1c).

Firstly, overlapping patches of size 512 × 512 pixels at 20× magnification were cropped from the tissue regions inside the slide in a sliding window fashion with stride 256 pixels. These patches were passed into the trained Mask R-CNN model to segment glands present.

Secondly, predicted masks from the Mask R-CNN model were gray-scale and converted to binary using thresholding. Then, binary masks were post-processed to merge partial predictions and eliminate redundant predictions arising from overlapping patch cropping (see Appendix B for details).

Finally, for each detected gland (instance) in a slide, patches at multiple resolutions were cropped from the slide. The trained multi-resolution model classified each detected gland as benign or malignant.

## 3 Results

### 3.1 Mask R-CNN model successfully segmented prostate glands

The Mask R-CNN model’s performance was evaluated on the test set of gland segmentation dataset. Using an intersection over union (IoU) threshold of 0.5, a recall of 0.945 and a precision of 0.830 were obtained at gland level. The low precision (compared to recall) was due to glands appearing inside dataset patches, but which were not annotated since they were partial glands at the edges of the biopsy cores (Figure 3b).

Moreover, we compared the Mask R-CNN model’s performance with other literature methods on a publicly available gland segmentation dataset [23] (Table 2). The Mask R-CNN model slightly outperformed deep learning-based segmentation methods. Besides, all the deep learning-based methods vastly outperformed traditional image processing or machine learning-based methods.

**Table 2:** Performances of different methods in prostate gland segmentation in terms of pixel-based metrics. The performances were on the test dataset of Salvi et al. [23]. Note that accuracy values were the balanced accuracy values as in Salvi et al. [23], and all the performance values except the one for the Mask R-CNN model were collected from Salvi et al. [23].

| Method | Accuracy | Precision | Recall | Dice |
| --- | --- | --- | --- | --- |
| Farjam et al. [24]<sup>*</sup> | $0.6378 \pm 0.1586$ | $0.7183 \pm 0.3034$ | $0.4372 \pm 0.1736$ | $0.5070 \pm 0.2059$ |
| Naik et al. [25]<sup>*</sup> | $0.7402 \pm 0.1151$ | $0.7958 \pm 0.2021$ | $0.5819 \pm 0.2275$ | $0.6357 \pm 0.2105$ |
| Peng et al. [26]<sup>*</sup> | $0.7957 \pm 0.1535$ | $0.6508 \pm 0.2568$ | $0.9305 \pm 0.1124$ | $0.7334 \pm 0.2198$ |
| Nguyen et al. [27]<sup>*</sup> | $0.7703 \pm 0.1632$ | $0.8260 \pm 0.1588$ | $0.7041 \pm 0.2998$ | $0.7145 \pm 0.2556$ |
| Singh et al. [28]<sup>*</sup> | $0.6734 \pm 0.1247$ | $0.9001 \pm 0.1743$ | $0.3869 \pm 0.2493$ | $0.4931 \pm 0.2557$ |
| Ren et al. [29]<sup>†</sup> | $0.8576 \pm 0.1139$ | $0.8199 \pm 0.1638$ | $0.8861 \pm 0.1673$ | $0.8308 \pm 0.1495$ |
| Xu et al. [30]<sup>†</sup> | $0.8250 \pm 0.1106$ | $0.7407 \pm 0.1597$ | $0.9273 \pm 0.1079$ | $0.8079 \pm 0.1264$ |
| Salvi et al. [23]<sup>†</sup> | $0.9325 \pm 0.0684$ | $0.8897 \pm 0.1359$ | $0.9356 \pm 0.0964$ | $0.9016 \pm 0.1087$ |
| Mask R-CNN [16]<sup>†,‡</sup> | **$0.9410 \pm 0.0010$** | **$0.9002 \pm 0.0026$** | **$0.9468 \pm 0.0011$** | **$0.9229 \pm 0.0015$** |

<sup>\*</sup> Traditional image processing or machine learning-based methods. <sup>†</sup> Deep learning-based methods. <sup>‡</sup> Standard deviations were calculated using bootstrapping [20].

### 3.2 The four-resolution model outperformed single-resolution models in gland classification

The four-resolution deep neural network model incorporated information from different levels in gland classification task. While 40× and 20× patches provided detailed morphology of the gland under consideration, 10× and 5× patches provided spatial neighboring information. We also trained single-resolution models for comparison. The models were evaluated using area under receiver operating characteristics curve (AUROC) and average precision (AP) calculated over precision vs. recall curve. 95% confidence intervals (CI) were constructed using the percentile bootstrap method [20].

The four-resolution model achieved the AUROC of 0.996 (95% CI: 0.994 - 0.997) and the AP of 0.997 (95% CI: 0.994 - 0.998). While the single-resolution models also produced satisfactory results, the four-resolution model outperformed them (Figure 2).

**Figure 2:**
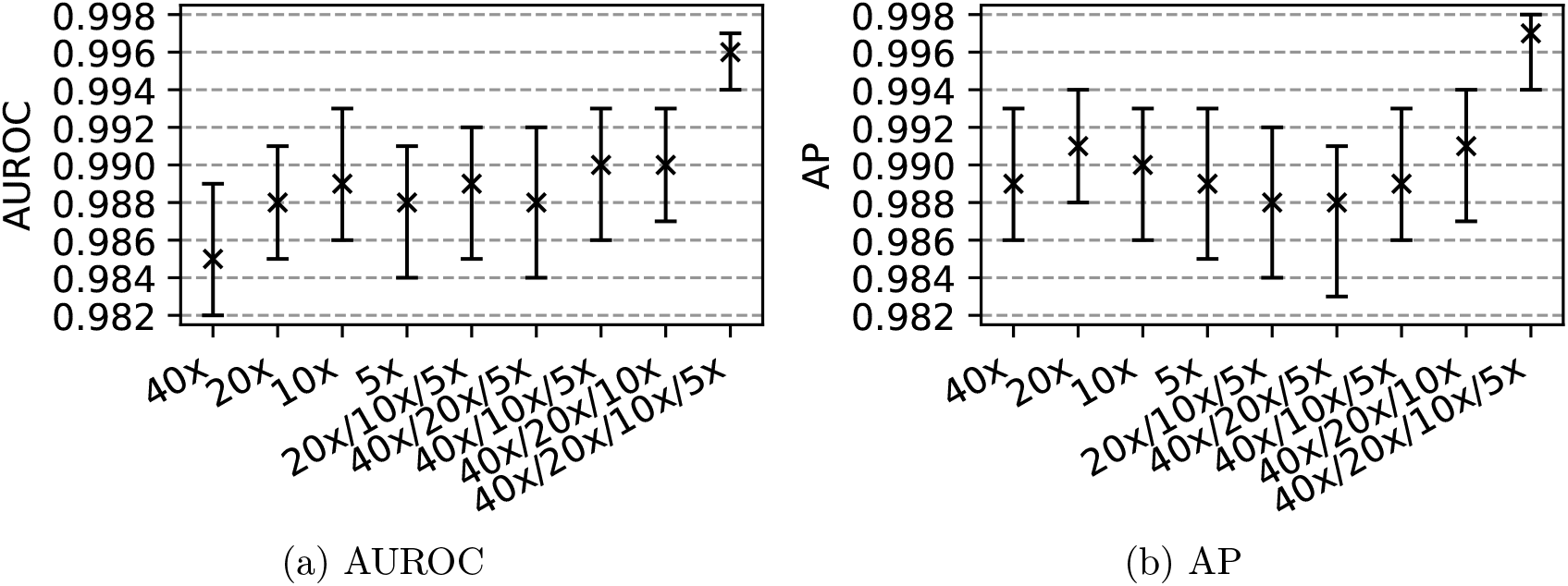
Performance evaluation on the test set of gland classification dataset. (a) Area under receiver operating characteristics curve (AUROC) and (b) average precision (AP) calculated over precision vs. recall curve together with 95% confidence intervals (obtained using the percentile bootstrap method [20]) are presented for the single resolution models, three-resolution models, and four-resolution model.

### 3.3 Gland morphology and neighborhood info were important in prostate gland classification

To assess the contribution of each resolution to the gland classification performance, we trained three-resolution models by dropping a different resolution each time from the four-resolution model. Then, the performance drop on the test set was used as a metric for that particular resolution. The highest and second-highest performance drops were observed in both AUROC and AP when the resolutions of 10× and 40×, respectively, were excluded (Figure 2). Our analysis showed that 10× and 40× patches provided valuable information for the four-resolution model. This also validated that both morphology (from 40×) and neighborhood (from 10×) info were important in prostate gland classification.

### 3.4 The pathologist’s annotations served as landmarks and guided the researcher in creating viable annotations for machine learning

Asking a pathologist to annotate every single gland in a slide is expensive and not feasible. Therefore, we followed a different strategy (see Appendix A for details). A senior pathologist annotated only 10% of the glands in each slide. Based on these annotations, a researcher annotated the rest of the glands.

We checked the effectiveness of our annotation strategy. In the test set of gland classification dataset, the four-resolution model’s performance was calculated on only the glands annotated by the pathologist and only the glands annotated by the researcher. The AUROC of 0.995 (95% CI: 0.991 - 0.999) and the AP of 0.996 (95% CI: 0.992 - 0.999) were obtained on the glands annotated by the pathologist. Similarly, the AUROC of 0.996 (95% CI: 0.993 - 0.997) and the AP of 0.997 (95% CI: 0.994 - 0.998) were obtained on the glands annotated by the researcher. Obtaining similar performance on both subsets of the test set, we concluded that the pathologist’s annotations served as landmarks and guided the researcher in creating viable annotations for machine learning.

### 3.5 Deep learning-based pipeline successfully classified biopsy parts into negative and positive

There were multiple needle biopsy cores in a WSI, and these cores could be broken into parts during slide preparation. Each core needle biopsy part within WSIs in the test set of gland classification dataset was classified into positive vs. negative based on the manual annotations. A part was assigned a positive label if it contained at least one malignant gland and a negative label otherwise. Then, the pipeline was tested end-to-end on the core needle biopsy part classification task (81 parts in 16 slides of 16 patients in the test set: 50 benign and 31 malignant). The glands in each part were detected by the trained Mask R-CNN model. For each detected gland, a malignancy probability was obtained from the trained four-resolution model (see Methods 2.4). The maximum of the predicted malignancy probabilities in a part was used as the part’s malignancy probability. An AUROC value of 0.997 (95% CI: 0.987 - 1.000) was obtained (Table 3).

**Table 3:** Performance of algorithms in prostate cancer detection in core needle biopsy slides.

| Study | Internal Test Dataset (benign:malignant) | AUROC (95% CI) |
| --- | --- | --- |
| Campanella et al. [6] | 1274 core needle biopsy slides (1439 benign, 345 malignant) | 0.991 (0.986 - 0.995)* |
| Pantanowitz et al. [10] | 2501 core needle biopsy slides (1957 benign, 490 malignant) | 0.997 (0.995 - 0.998) |
| Ström et al. [11] | 1631 core needle biopsy slides (910 benign, 721 malignant) | 0.997 (0.994 - 0.999) |
| Bulten et al. [12] | 535 core needle biopsies (250 benign, 285 malignant) | 0.990 (0.982 - 0.996) |
| Current study | 81 core needle biopsy parts ( 50 benign, 31 malignant) | 0.997 (0.987 - 1.000) |

AUROC: area under receiver operating characteristics curve, CI: confidence interval.
\*CI was obtained on provided predictions in the paper using the percentile bootstrap method [20].

Furthermore, to assist pathologists in reading prostate core needle biopsy slides, a spatial malignancy map for each slide was constructed using malignancy probabilities of detected glands obtained from the trained four-resolution model (Figure 3).

**Figure 3:**
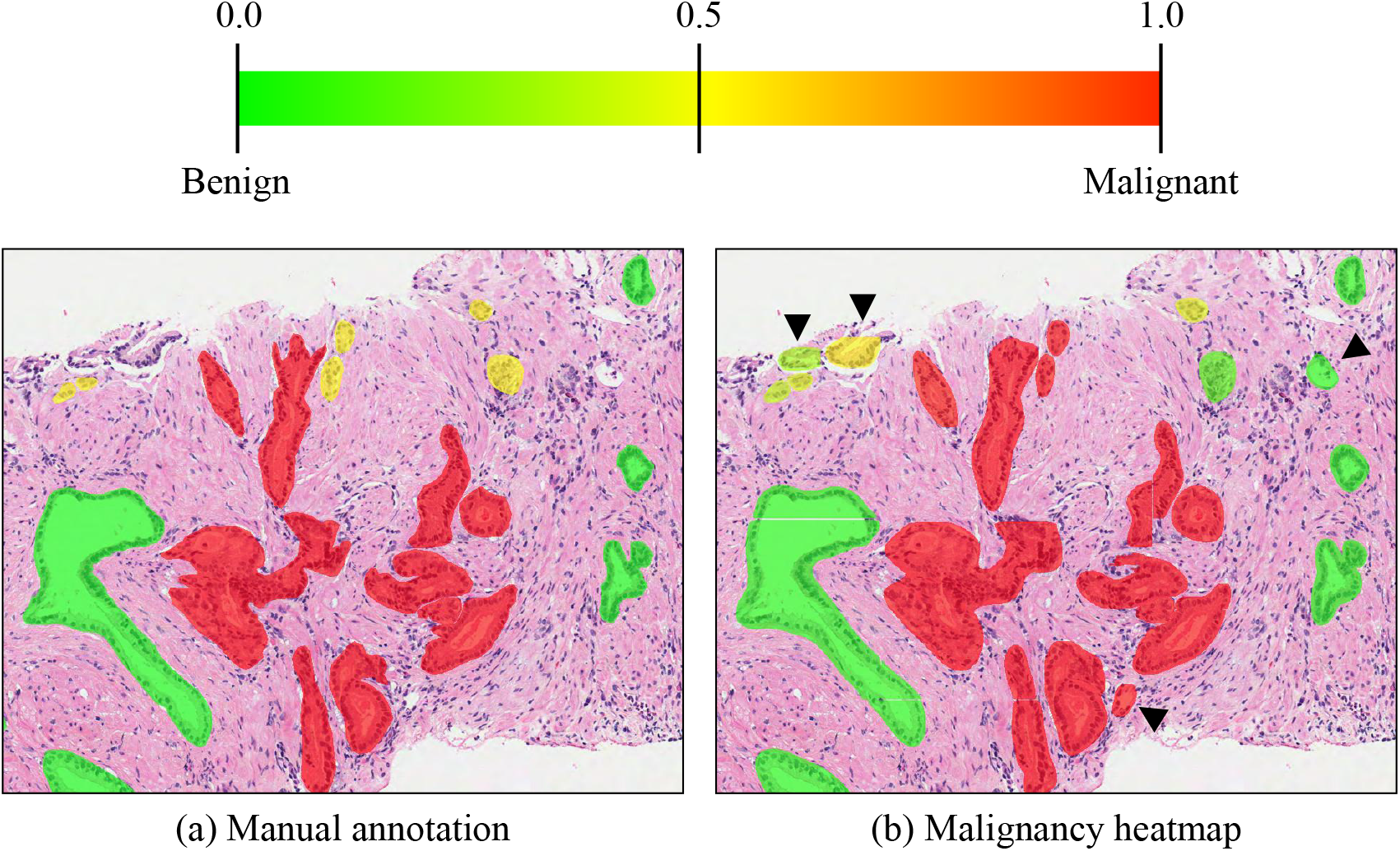
Spatial malignancy map. (a) and (b) show example manual annotations and spatial malignancy map produced using deep learning-based pipeline, respectively. Arrow heads show tissue components detected by the pipeline but not manually annotated.

## 4 Discussion

Manual reading of core needle biopsy slides by pathologists is the gold standard in the prostate cancer diagnosis in the clinic. However, it requires the analysis of around 12 (6-18) biopsy cores, including hundreds of glands. Especially for low-grade and low-volume prostate cancer (GS 3+3 and 3+4), identifying the few malignant glands among vastly benign glands is a tedious and challenging task. These few malignant glands can be easily overlooked, potentially resulting in missed therapeutic opportunities. This study developed a deep learning-based pipeline to detect malignant glands in core needle biopsy slides of low-grade prostate cancer patients. The pipeline can help early diagnosis of prostate cancer and effective use of other therapeutic tools at early stages, like active surveillance, rather than aggressive prostatectomy eligible for later stages. Moreover, the pipeline can reduce the pathologists’ workload as an assistive tool.

### 4.1 Deep learning-based pipeline can assist pathologists

Our pipeline successfully classified biopsy cores into negative and positive. It can be deployed as a pre-analysis stratification tool and help pathologists effectively manage their time on each biopsy core. For instance, they can spend less time validating negative cores while devoting more time to positive cores. Besides, spatial malignancy maps can help them concentrate on high-risk regions and decide if further cuts are required (for example, in the regions with a malignancy probability of 0.5) to make a diagnosis.

Furthermore, our pipeline can be deployed as a second-read system. The system can generate a flag for a second opinion in case of a contradiction between the pathologist’s diagnosis and the system’s classification. This can help reduce false negatives and false positives which potentially result in missed therapeutic opportunities and aggressive treatment, respectively.

### 4.2 Challenges of gland-level annotations

Gland-level manual annotation is a challenging task, especially in core needle biopsies. A tissue core starts drying out from the surface after excision. If the core is not put into formalin buffer immediately, the morphology of the glands at the edges becomes distorted, making benign vs. malignant classification more challenging. Besides, the glands at the edges are usually partial, and partial glands are not used for diagnosis in the clinical routine. If all the glands within the core are partial, pathologists usually make a diagnosis based on other viable cores. Moreover, most glandular structures inside the cores are tangential cuts, which are hard to annotate. They are considered secondary for diagnosis and mostly require deeper cuts to reveal the glandular structure for diagnosis.

Furthermore, artifacts occur during the sample preparation, such as detached glands, folded tissue, uneven cuts, and poor preservation. These also make the annotation of each gland challenging. Another challenge appears in identifying the boundaries of the glands. It can be difficult to draw the boundaries of branching glands and fused glands during manual annotation.

Despite these challenges, gland-level annotations provide a fine-level resolution to identify individual glands as benign or malignant. This helps us train machine learning models with fewer slides than we would need with slide-level annotations.

### 4.3 Limitations and Future Work

Gland-level annotations enabled us to train highly accurate machine learning models. However, we had a limited number of annotated slides since manual annotation was tedious and time-consuming. It would have been better if we had more slides to consolidate our model’s performance. Another limitation of our study was the lack of external validation. Although we tested our model’s performance on slides of hold-out patients never seen by the model, an external cohort study would have shown the robustness of our model against inter-institution differences.

In the future, we wish to deploy our pipeline as a second-read system in Singapore and check its performance in the real-world clinical flow.

## 5 Contributors

MUO, MYN, CECX, and LAYX selected the patients and collected the data. MYN and DMG annotated the data. MUO, MYN and MS verified the underlying data, conducted the experiments and analyzed the results. MUO and MYN wrote the manuscript. MS, WY, WKS, CFW and HKL contributed to the manuscript preparation. CFW and HKL supervised the study. All authors reviewed the manuscript and agreed with its contents.

## 7 Data sharing

All original code has been deposited at Zenodo under the https://doi.org/10.5281/zenodo.5982398 and publicly available. The repository provides a detailed step-by-step explanation, from training of gland segmentation and classification models to inference with the trained models. All the images have been deposited at Zenodo under the https://doi.org/10.5281/zenodo.5971764 and publicly available [19].

## 8 Acknowledgments

This study was funded by the Biomedical Research Council of the Agency for Science, Technology and Research, Singapore.

## Appendix A Annotation workflow

The workflow to annotate core needle biopsy slides is given in Figure 4.

**Figure 4:**
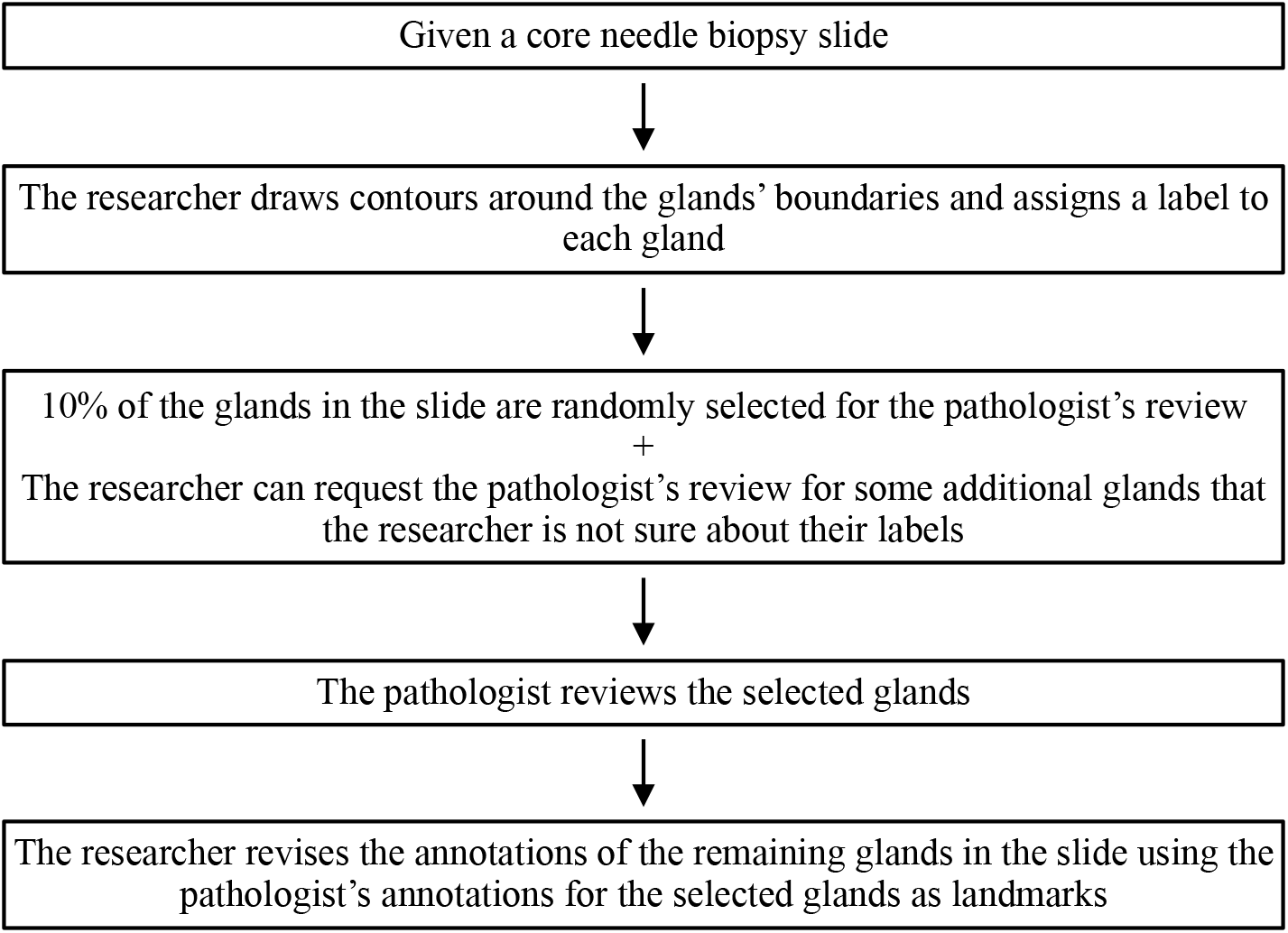
The workflow for the annotation of core needle biopsy slides.

## Appendix B Post-processing predicted masks

Predicted gray-scale mask for each instance was thresholded at 0.5 to obtain a binary mask. When thresholding resulted in multiple contours, the largest contour was used to represent a segmented instance (gland). Holes in the binary mask, if any, were then filled. Predictions at tissue boundaries extending to the background were excluded.

Multiple binary masks corresponding to the same gland were observed due to the use of overlapping patches and detection of incomplete glands at the boundary. Thus, the masks were processed to remove redundant ones as follows:

1. When the intersection-over-union (IoU) or intersection over minimum area between two predicted masks exceeded the threshold of 0.3, the mask with the lower prediction score was discarded.
2. A mask that intersected with two or more masks was excluded.
3. Finally, two binary masks that had an IoU exceeding 0.3 were merged in an iterative manner starting with the pair of masks that had the greatest IoU.

